# Aging diminishes interlaminar functional connectivity in the mouse cortical V1 and CA1 hippocampal regions

**DOI:** 10.1101/2025.09.17.676336

**Authors:** Teresa Thai, Steven M. Wellman, Naofumi Suematsu, Camila Garcia Padilla, Te-Hsuan Tung, Sadhana Sridhar, Franca Cambi, Takashi D.Y. Kozai

## Abstract

Aging disrupts brain network integration and is a significant risk factor for cognitive decline and neurological diseases, yet the circuit-level mechanisms underlying these changes remain unclear. Most previous studies have utilized cross-sectional or acute approaches, limiting insights into the longitudinal dynamics of the neural network. In this study, we chronically recorded laminar electrophysiological activity in both the primary visual cortex (V1) and hippocampal CA1 region of young (2-month-old) and aged (13-month-old) mice over 16 weeks. This approach allowed us to directly assess how aging modulates functional connectivity within hierarchically connected cortical and hippocampal circuits. We found that single-unit spiking activity and the signal-to-noise ratio were largely preserved in aged versus young mice, suggesting intact neuronal firing properties. However, aged mice showed global reductions in local field potential (LFP) power and a selective decrease in coherence across delta, alpha-beta, and gamma frequency bands within and between cortical layers and V1-CA1 pathways, while phase amplitude coupling remained unaffected. Interestingly, population level excitatory activity in CA1 was increased in aged animals. These findings indicate that aging selectively impairs network-level synchrony and temporal coordination in specific frequency bands and regions, with minimal loss of single-neuron function. Our results highlight the necessity of longitudinal, multi-region measurements to uncover the multi-scale vulnerabilities of the aging brain. Understanding the depth- and region-dependent circuit changes will guide strategies to preserve cortical-hippocampal communication and cognitive function in aging, as well as enhance neural interface technologies for older populations.

**NEW & NOTEWORTHY:** Aging non-uniformly degrades cortico-hippocampal networks, leading to region- and frequency-specific breakdowns in coordinated activity. While single-neuron activity remains preserved, disruptions in frequency-selective synchrony (delta and alpha-beta) were observed in aged mice, indicating impaired V1-CA1 communication as a marker of aging rather than neuronal degeneration. By separating single-neuron activity and large-scale dynamics, we reveal that aging alters communication between sensory and memory systems, underscoring the need for longitudinal approaches to capture age-related impairments in laminar connectivity.

## INTRODUCTION

Aging is a significant risk factor for a wide array of neurological disorders, such as Alzheimer’s disease (AD), and is associated with substantial cognitive decline in a major segment of the population (Haberman et al., 2017; Hou et al., 2019; Voytek & Knight, 2015). This decline manifests across various cognitive domains, including memory, processing speed, and executive functions, affecting daily tasks and quality of life (Zhang, 2023). Beyond functional changes, the aging brain undergoes significant structural alterations, such as cerebral tissue volume loss (atrophy), cortical thinning, sulcal widening, and ventricular enlargement (Blinkouskaya et al., 2021). The trajectory of these changes is highly heterogeneous across individuals, with variations in molecular, cellular, and phenotypic changes due to diverse genetic susceptibility and environmental exposures.

### Age-driven deficits in cortical and hippocampal laminar circuits

A key approach to understanding age-related brain changes involves examining functional and structural connectivity, providing insights into the brain’s intrinsic organization and information flow (Chen et al., 2024; Damoiseaux, 2017). On a larger scale, healthy aging is often associated with reduced global efficiency, characterized by weakened long-range connections and higher functional segregation (Ajilore et al., 2014; Iordan et al., 2017). While within-network functional connectivity often decreases, some studies indicate increased between-network functional connectivity in older adults (Du et al., 2023; Jauny et al., 2024), which may represent a compensatory mechanism. These network alterations are observed across multiple brain regions, including sensorimotor, visual, frontal-occipital, and cerebellar systems (Filippi et al., 2023; Shah et al., 2018). Structural connectivity studies also show an inverted U-shape trajectory for local and global efficiency with age (Khalilian et al., 2024; Niu et al., 2022), indicating lower network efficiency and altered brain dynamics. However, much of this understanding comes from cross-sectional human studies (Biswal et al., 1995; Watanabe et al., 2021), emphasizing the critical need for longitudinal animal studies to track the evolution and causality of these changes and to determine the order of age-related changes, such as whether changes in brain structure precede functional or cognitive changes. Such longitudinal designs are essential for establishing causal relationships and enabling early detection and targeted interventions. While these macroscale network changes provide critical insight, how aging influences finer circuit-level processes that underline large-scale dynamics remain elusive.

On a finer scale, interlaminar connectivity coordinates information transfer between cortical layers and has emerged as a critical node for pinpointing age-related disruptions. These vertical interactions allow feedforward and feedback communication to be integrated across layers and between brain regions.

This study focuses on the primary visual cortex (V1) and hippocampal CA1, key regions for visual processing and memory integration, respectively. Disruptions at either site may fragment the broader communication axis and weaken the V1-CA1 functional connectivity (Preston et al., 2025; Tran et al., 2016). As V1 and CA1 regions are architecturally distinct, assessing their differences proves a window into how aging alters both cortical laminar processing and hippocampal integration. Within the V1-CA1 communication axis, age-related changes affect the precise timing synchrony of neural oscillations (Tran et al., 2020; Voytek et al., 2015). Previous work in young mice has shown depth-dependent degradation in intralaminar functional connectivity near implanted microelectrodes, with early impairment noted in superficial layers (LII/III) and specific directional changes in interlaminar functional connectivity between LII/III and LV (Chen et al., 2024). In the hippocampal CA1 region, changes in the precise timing of spikes in response to oscillatory local field potential (LFP) activity have also been observed (Newman et al., 2017), with alterations in locomotor behavior and spatial memory (Crown et al., 2022). These findings illustrate that laminar-specific impairments are not isolated circuit defects, but rather disruptions with far-reaching consequences of functional connectivity. These specific localized impairments highlight the need to move beyond macroscale observations to unravel the underlying cellular and molecular mechanisms of age-related neurological dysfunction, particularly in relation to the brain’s response to chronic implants.

### Intersection of aging and implant-induced impairments

Although age-related vulnerabilities contribute to the decline in neural signal quality and functional connectivity, these effects are further amplified by chronic implantation. During normal aging, some cortical areas exhibit enhanced myelination, pointing to oligodendrocytes as a potential glial cell of interest for reversing age-related degradations (Hill et al., 2018; Murray et al., 2023); while other regions experience atrophy, demyelination, and cognitive deficits (Fujita et al., 2023). As myelin and axonal integrity are critical for saltatory conduction, delays and asynchronous signaling across brain regions degrades the functional connectivity and efficiency of large-scale networks. Aging also reduces neurovascular coupling, increases oxidative stress, and promotes neuroinflammation (Liguori et al., 2018; Yabluchanskiy et al., 2020), limiting the ability of neural populations to maintain cortico-hippocampal synchrony. These impairments interact with disrupted autophagy and lysosomal function, resulting in lipofuscin accumulation and misfolded protein deposition (amyloid-beta and tau) as well as compromising synaptic plasticity. Notably, these cellular processes are exacerbated by chronic implantation (Chen, Padilla, et al., 2023; Wellman et al., 2023), suggesting a potential physiological link between aging and implant-induced neurodegeneration. Senescent cells release pro-inflammatory SASP factors (Shafqat et al., 2023) and reactive glial cells proximal to implants decline in function (Chen et al., 2024; Salatino et al., 2017; Wellman et al., 2018), leading to impaired blood-brain barrier integrity that is exacerbated by age and APOE4 status (Nelke et al., 2022). Finally, receptor alterations, including age-dependent reductions in NR2B subunits of NMDA receptors, may disrupt synaptic plasticity critical for memory (Gaikwad et al., 2024). Together, these interconnected mechanisms emphasize how aging and chronic implantation interact to accelerate neurodegeneration, reinforcing the need to assess longitudinal network-level consequences.

Despite the growing understanding of age-related brain changes at various scales, there remains a critical gap in longitudinally characterizing how these complex multi-scale alterations directly impact the functional neural network connectivity in the living brain over time, particularly at the microcircuit level. While many studies rely on cross-sectional data or macroscale imaging, it remains unclear whether modulations in network dynamics originate from single-neuron alterations or population-level dysfunction. This study directly addresses these limitations by employing chronic, multi-site electrophysiological recordings from key brain regions, primary visual cortex (V1) and hippocampal CA1, in young (2-month-old) and aged (13-month-old) C57BL6 wild type mice over a 16-week period. Single-unit and multi-unit metrics, including yield and signal-to-noise ratio, were largely preserved with aging, indicating that individual neuron activity remains intact. In contrast, aged mice exhibited reduced local field potential (LFP) power and decreased Δcoherence across delta, alpha-beta, and gamma bands between cortical layers and CA1, while phase-amplitude coupling remained stable. These findings demonstrate that aging selectively impairs network-level connectivity and temporal coordination of hierarchical pathways without substantially altering single-neuron firing. By revealing these multi-scale disruptions, this work highlights the importance of laminar and interregional network integrity in maintaining cognitive function and has direct relevance for older populations receiving brain implants, where age-dependent network vulnerabilities and altered neurovascular responses may affect implant performance and longevity.

## MATERIALS AND METHODS

### Animals

All use of animals in the present study followed the guidelines of Institutional Animal Care and Use Committees (IACUC) of the Veterans Administration Pittsburgh Health Sciences (Animal Welfare Assurance number A3376-01) and of the University of Pittsburgh (Animal Welfare Assurance number A3187-01). Protocols (#0321 and #15043715) were approved by IACUC. All mice were bred and housed in a barrier facility with a 12-hour light/dark cycle. Food and water were available ad libitum.

### Surgical Microelectrode Implantation

Prior to microelectrode implantation, wild-type mice (C57BL/6J) were aged to 2 months (young, N = 6 males) or 13 months (aged, N = 2 males, 4 females). The young group included 6 male mice, while the aged group consisted of 2 male and 4 female mice. In this study, single-shank Michigan-style electrodes (NeuroNexus, A1×16-3mm-100-703-CMLP16) were perpendicularly implanted into the left priminary monocular visual cortex (V1) using previously published methods (Chen, Cambi, et al., 2023b; Chen et al., 2024; Steven M. Wellman et al., 2020). Probes were implanted at a depth of 1600μm, spanning the entire cortical column and the superficial layers of the CA1 subfield of the hippocampus. Briefly, animals were anesthetized with a ketamine (75 mg/kg) and xylazine (7 mg/kg) cocktail and injected intraperitoneally (I.P.) prior to fixing the animal onto a stereotaxic frame. Body temperature was monitored and maintained using a rectangular heating pad (Deltaphase isothermal pad, Braintree Scientific Inc. Braintree, MA). Hair, skin and connective tissue were removed to expose the top of the skull. Vetbond adhesive was applied to the top of the skull to dry the surface prior to drilling. Two bone screw holes were drilled over the motor cortices, and one was drilled over the contralateral visual cortex. Stainless steel bone screws that were 4 mm long and 0.86 mm in diameter (Fine Science Tools, British Columbia, Canada) were positioned into the holes to connect the electrode’s ground and reference wires as well as anchoring the headcap. A craniotomy 1mm anterior to Lambda and 1.5 mm lateral to the midline was performed using a high-speed dental drill and 0.7 mm drill bit. Saline was periodically applied to the surgical site to prevent thermal damage during the drilling process. A microelectrode was then inserted at a speed of 15 mm/s with a DC motor-controller (C-863, Physik Instrutmente, Karlsruhe, Germany). The last channel site was visually confirmed to penetrate below the pial surface to target CA1 subregion of the hippocampus. A silicone elastomer (Kwik-Sil) was used to fill the craniotomy site and space between the exposed brain and microelectrode shank. Dental cement cap was performed to seal the surgical site. Mice were given an I.P. injection of ketofen (5 mg/kg) on the day of surgery and two days post-operation.

### Electrophysiological Recordings

Electrophysiological recordings were performed inside a grounded Faraday cage to minimize electrical and environmental noise as previously described (Alba et al., 2015; Kolarcik et al., 2015; Kozai et al., 2016; Kozai et al., 2015; Nicolai et al., 2018). Mice were placed on a rotating stage for awake, head-fixed recordings in a dark room. For visually evoked neural activity, a MATLAB-based Psychophysics toolbox was used to display a drifting gradient on a 24” LCD monitor (V243H, Acer, Xizhi, New Taipei City, Taiwan) approximately 20 cm away from the eye contralateral to the implanted hemisphere, covering a 60° wide by 60° high visual field. The drifting gradient of solid white and black bars were displayed and synchronized with the electrophysiology recording system (RX7, Tucker-Davis Technologies, Alachua, FL) at a sampling rate of 24,414 Hz. The white and black grating stimulus was presented for 1 s and rotated at 135° increments followed by 1 s dark screen. Each recording session had a total of 64 stimulation trials across a 16-week period, measuring daily activity within the first 14 days followed by weekly recordings thereafter.

### Current Source Density

To account for any drifting and brain swelling within the initial days post-implant, we used the current source density (CSD) during evoked activity in the visual cortex to identify layer IV along the length of the probe. The CSD plots were generated by averaging the evoked local field potential (LFP) for each electrode site, smoothed the signal across all 16 sites, and then calculated the second spatial derivative. The CSDs were then averaged over 64 stimulus trials. Layer IV was identified by the inversion of LFP polarity occurring within the first 100 ms post-stimulus onset, monitoring changes in Layer IV position over the 16-week experimental period. The amount of cortical drift over time was reported as the mean change in layer IV depth relative to the depth identified at day 0. All electrophysiological analyses along cortical depth were performed after aligning electrode sites to the corresponding layer IV depth for all animals in each group and for each day. The hippocampal region was identified as the electrode site beneath the electrically silent white matter tracts.

### Single-unit (SU) Analysis

A custom MATLAB script was used to process the raw signal data offline modified from previously established techniques (Kozai et al., 2015). Raw data was processed using a band-pass Butterworth filter from 2 to 300Hz to isolate LFPs and 300 Hz to 5kHz for neuronal spiking activity. Common average referencing was applied to the data stream as described (Ludwig et al., 2009). A threshold of 3.5 standard deviations below the mean was applied to distinguish potential neuronal single-unit (SU).

Channels with peak-to-peak voltage >10 μV and signal-to-noise ratio (SNR) >2 and were considered for single-unit sorting. Active sites were defined as any electrode channel with discernable SUs. SNR was calculated as the peak-to-peak voltage for each single unit divided by the noise and reported as the average SNR and average SNR per active site. Analyzing the mean waveform shape, Signal-to-noise ratios (SNR) were calculated by dividing the peak-to-peak amplitude of each single unit by the noise and reported as average SNR and average SNR per active site (electrode channels reporting detection of SU) over time. Sortable single units were confirmed by observing the quality and shape of neuronal waveforms, auto-correlograms, and peri-stimulus time histograms (PSTH) with 50ms bins. SU yield was calculated as the percentage of electrode sites (out of 16) with at least one identifiable single unit. The noise floor for each electrode site was taken as two times the standard deviation (2*STD) of the entire recorded data stream after extracting all threshold crossing events. Finally, the SNR of any channels without a sortable SU waveform were reported as zero when calculating averages.

### Multi-unit (MU) Analysis

Multi-unit (MU) activity was defined as any recorded threshold crossing event that occurred within 1s after the onset of the visual stimulus trigger or pseudotrigger. Peri-stimulus time histograms (PSTHs) of 50ms bins were used to assess the dynamics of MU activity. The firing rate was calculated as the mean number of threshold crossing events in a 1s period after the onset of the stimulus trigger or pseudotrigger. Evoked MU activity was assessed by calculating multi-unit yield and signal-to-noise firing rate ratio (SNFRR). Parameters for MU spike count were determined by varying the temporal bin size and post-stimulus latency, ranging from 0 to 1s in 1ms increments. MU yield was defined as the number of electrode sites that exhibited a statistically significant difference (paired t-test, p < 0.05) in spike count for a given bin size and latency after stimulus onset (stim ON) compared to the corresponding pre-stimulus interval (stim OFF). SNFRR evaluates the change in MU firing rate before and after stimulus presentation, normalized by the mean standard deviation across conditions:

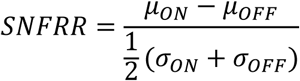

Across 64 trials, *μ*_*ON*_ and *μ*_*OFF*_ represent the mean firing rates during stimulus ON and OFF conditions, and *σ*_*ON*_ and *σ*_*OFF*_ are the standard deviations of firing rates during ON and OFF conditions, respectively. A higher SNFRR value indicates elevated visually evoked firing activity. We reported the magnitude of modulation of visually evoked neural activity, |SNFRR|.

### Local Field Potential Analysis

LFP power spectra were calculated using a multi-taper method of 1 s duration, 1 Hz bandwidth, and a taper number of 1. Relative power was calculated as the ratio of power within a specific frequency band to the total LFP power across the entire frequency range. To normalize LFP power, the spontaneous power spectrum was subtracted from the evoked power spectrum. Relative power and the normalized power spectrum were calculated as follows:

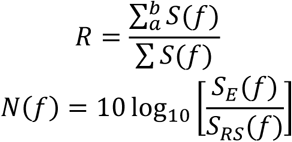

Here, *R* represents the relative power, *S*(*f*) is the LFP power spectrum, *α* and *b* are the lower and upper limits of the specified frequency range, *N*(*f*) is the normalized power spectrum, and *S*_*E*_(*f*) and *S*_*RS*_(*f*) represent the evoked and resting-state power spectra.

### Laminar Coherence Analysis

The intra- and inter-laminar depths was calculated as the magnitude-squared coherence as previously described (Michelson & Kozai, 2018). Coherence quantitatively describes the similarity between two signals based on their frequency-dependent responses. The coherence was reported as values ranging from 0 to 1, indicating no relationship or a perfect linear relationship between two corresponding signals, respectively. Coherence values were calculated using the following equation:

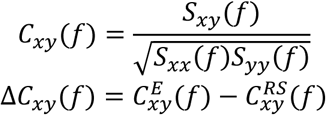

Here, *C*_*xy*_(*f*) represents coherence, *S*_*xy*_(*f*) is the cross spectrum, and *S*_*xx*_(*f*) and *S*_*yy*_(*f*) are the autospectra of LFP activity between electrode sites *x* and *y*, respectively. Following this, the data were normalized by subtracting the resting-state coherence 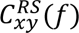 for the evoked coherence 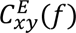 to give Δ*C*_*xy*_(*f*). Calculations were done within 1 s intervals after each stimulus presentation or pseudotrigger. A 3 Hz half-bandwidth and taper number of 5 was used before averaging across all 64 trials. The average coherence of all electrode sites were compared between groups and different brain regions. The coherence values for each frequency band were reported as the average coherence within that frequency range across all animals per group. Thus, the range of Δcoherence is −1.0 to 1.0, where negative values represent higher synchronicity during the resting state and positive values represent higher synchronicity during the evoked state.

### Phase Amplitude Coupling

To quantify the directionality of neuronal circuit connections, the phase amplitude coupling (PAC) was calculated based on Kullback-Leibler (KL) formula as previously described (Chen et al., 2024; Chen, Padilla, et al., 2023; Scheffer-Teixeira & Tort, 2016; Tort et al., 2010). The raw signal was processed with a bandpass filtered for specific LFP frequencies *f*. The Hilbert transform was applied to output the time series of the phase component from the slow frequency LFP activity in Channel X, denoted as *ϕ*_*x*_(*t*, *f*_*x*_) where *t* represents time and *f*_*x*_ represents phase frequencies 2,3,4… 20Hz. The time series of the amplitude component at high frequency LFP activity in channel Y (*A*_*Y*_(*t*, *f*_*Y*_)) was extracted using Morlet wavelet transform (5 cycles) per frequency, using amplitude frequencies *f*_*Y*_ = 30,31,32…90Hz.

Next, we calculated the amplitude of high frequency LFP oscillation in Channel Y at each given slow frequency LFP activity in Channel X, resulting in the composite time series *ϕ*_*x*_(*t*, *f*_*x*_), *A*_*Y*_(*t*, *f*_*Y*_). The phases were binned every 18° (N = total number of phase bins, 20) and the amplitude at each phase bin (*i*) was averaged to give *A*_*Y*_(*t*, *f*_*Y*_)_*ϕx*(*t*,*fx*)_(*i*). Lastly, normalizing by the sum of all bins, resulted in the normalized amplitude distribution over phases *P*(*i*, *f*_*x*_, *f*_*Y*_).

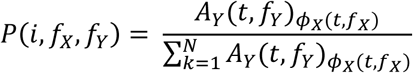

If there is no or minimal PAC between channels X and Y, the normalized amplitude distribution *P*(*i*, *f*_*x*_, *f*_*Y*_) would result in a uniform distribution. Further, the joint entropy defined as *H*(*f*_*x*_, *f*_*Y*_) measures the amount of deviation of the uniform distribution given by the following:

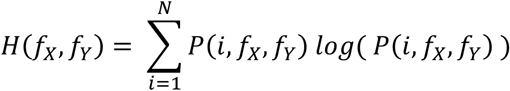

At the maximum join entropy *H*_*o*_ = *logN* ^2^, the normalized amplitude distribution *P*(*i*, *f*_*x*_, *f*_*Y*_) is uniform. We then calculated the difference between *H*(*f*_*x*_, *f*_*Y*_) and *H*_*o*_ using the KL distance formula. The modulation index (MI) of PAC was then calculated by dividing the KL distance by the uniform distribution *H*_*o*_.

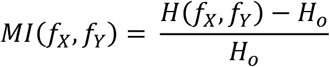

When Channel X = Channel Y, a higher MI value of PAC indicates stronger LFP synchronization coupling between the low frequency phase and high frequency amplitude and increased activation level of the intralaminar network. When Channel X ≠ Channel Y, a larger MI value of PAC suggests enhanced LFP synchronization coupling between the low frequency phase in the upstream layer and the high frequency amplitude in the downstream layer, resulting in an increase in the directional interlaminar circuitry.

### Electrochemical Impedance Spectroscopy

Impedance measurements were performed prior to each recording session using an Autolab potentiostat and a 16-channel multiplexer in awake, head-fixed mice. Measurements were performed using a 10 mV sine wave ranging from 10 Hz to 32kHz. The average impedance at 1 kHz was reported over all animals for each timepoint.

### Statistics

For metrics assessing changes in recording metrics over time, we fit the data to a linear mixed effects model. To model nonlinear relationships, a restricted cubic spline was applied with four knots placed at the 5th, 35th, 65th, and 95th percentiles of the data (Eles et al., 2017). Fixed effects included group (young and aged) and group-by-time interactions. Model significance was evaluated using a likelihood ratio test. Confidence intervals were calculated via bootstrapping with 1,000 iterations, with the 95% confidence interval defined as ± 1.96 times the standard error of the model estimates. Pairwise comparisons were further tested with post-hoc t-tests, applying a Bonferroni correction to control for type I error due to multiple comparisons.

Describe techniques, cell/animal models used (including species, strain, and sex), and lists of reagents, chemicals, and equipment, as well as the names of manufacturers and suppliers, including **<u>Research</u> <u>Resource Identifiers (RRIDs)</u>** when possible. Also in this section, describe the statistical methods that were used to evaluate the data. See **<u>Rigor and Reproducibility Guidelines (PDF)</u>** for more information.

All animal or human studies must contain an explicit statement that the protocols were reviewed and approved by an institutional review board or committee (<u>IACUC for example</u>), or that research was conducted under a valid license or permit from such a committee, board, or governing office (see **<u>Guiding Principles for the Care and Use of Vertebrate Animals in Research and Training</u>**). For human participants, written informed consent must be obtained. If not, provide an explanation. If clinical trials were used, a statement of registration and the clinical trial number are required.

## RESULTS

### Aging does not impact recording quality of single-unit neural activity

To assess the effect of aging on electrophysiological properties of single neurons, extracellular recordings were obtained using 16-channel intracortical microelectrode arrays implanted 1.6mm in 2-month-old or 13-month-old wild-type (WT) animals, targeting both the primary visual cortex (V1) and CA1 subregion of the hippocampus (Figure 1A). Assessing over depth, aged animals had increased activity yield in superficial regions with diminishing signal detection in deeper areas over time compared to young animals (Figure 1B). However, quantifying this trend, we observed no differences in the mean single-unit (SU) yield and mean signal-to-noise ratio (SNR) between aged and young mice across the 16 weeks (Figure 1C-D). There was a notable decline in electrophysiological measurements in both groups across time, consistent with previously published studies utilizing a similar microelectrode design (Chen, Cambi, et al., 2023; Kozai et al., 2014; Sturgill et al., 2022; Steven M Wellman et al., 2020). To account for possible glial scarring and white matter tracts separating V1 and CA1 that are devoid of neurons, we evaluated the mean SNR per active site, defined as electrode sites that had a discernable unit. We found that mean SNR/active site was comparable between young and aged animals (Figure 1E) with an interaction effect between time and group (p = 1.68e-07, likelihood ratio test). This may indicate that signal detection is more stable in aged animals over time following microelectrode implantation. Lastly, we assessed changes in electrochemical impedance spectroscopy as an indicator of microelectrode integration into brain tissue over the 16-week period. Although the young mice had slightly higher impedance, possibly an indicator of elevated glial response (Biran et al., 2005; Jeakle et al., 2023; Wellman et al., 2018), the values were comparable between young and aged mice (Figure 1F). Together these results indicate that aging does not impair neural signal quality or electrode performance, with increased stability for chronic microelectrode-tissue integration.

**Figure 1.**
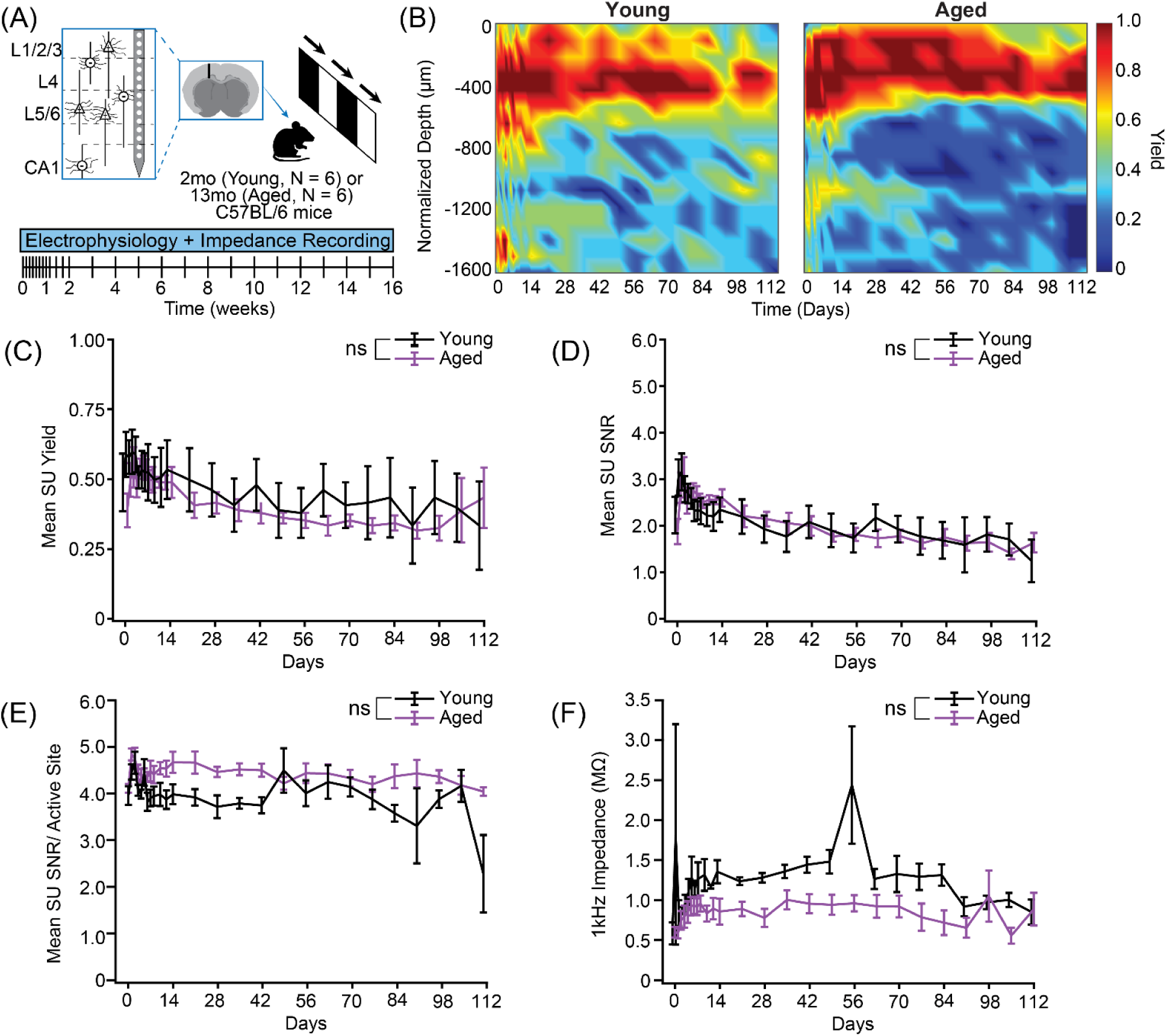
Aging does not impact single-unit detectability and activity. (A) Graphical representation of experimental model and timeline used for longitudinal electrophysiological recording (16 weeks). C57BL/6 wildtype (WT) animals at 2 months (young, N = 6 male mice) or 13 months (aged, N = 2 male, 4 female mice) were implanted with a 16-channel microelectrode array, spanning all cortical layers of V1 and CA1 subregion of the hippocampus (1.6 mm). (B) Heatmaps across time and normalized electrode implantation depth of signal yield for young and aged animals shows laminar differences in electrode channel yield. (C-F) Single-unit (SU) electrophysiological metrics over time for young (black solid) versus aged mice (purple solid) represented as mean ± SEM per timepoint (C) Mean SU yield over time and (D) mean SU signal-to-noise ratio (SNR) were comparable between young and aged groups. (E) Mean SU SNR per active site was slightly elevated in aged animals but insignificant. (F) Average impedance at 1 kHz over time. Non-significant differences are indicated by ns, determined by linear mixed effects model between young and aged mice.

### Putative excitatory activity is elevated in CA1 but not cortical layers of V1

To begin assessing differences in V1 cortical regions and CA1 subregion of the hippocampus, we aligned electrophysiological recordings to layer IV, identified by aligning sensory-evoked local field potentials with current source density as aforementioned (Figure 2A) (Chen, Cambi, et al., 2023; Chen et al., 2024). As modulations in excitatory-inhibitory circuits can contribute to measurements of signal detectability and spiking activity, we putatively labeled inhibitory and excitatory units based on the trough-to-peak interval (TPI) of the waveform shape. A Gaussian mixture model was fit to the TPI distribution, defining a threshold of 0.51ms. Units with TPI < 0.51 ms were putatively classified as inhibitory units and TPI ≤ 0.51 ms were putatively classified as excitatory units (Figure 2B). Representative waveforms were included in the inset of the TPI histogram. In the cortex, we found that, on average, the decline of signal detection of inhibitory and excitatory activity was comparable between young and aged animals (Figure 2C-D).

**Figure 2.**
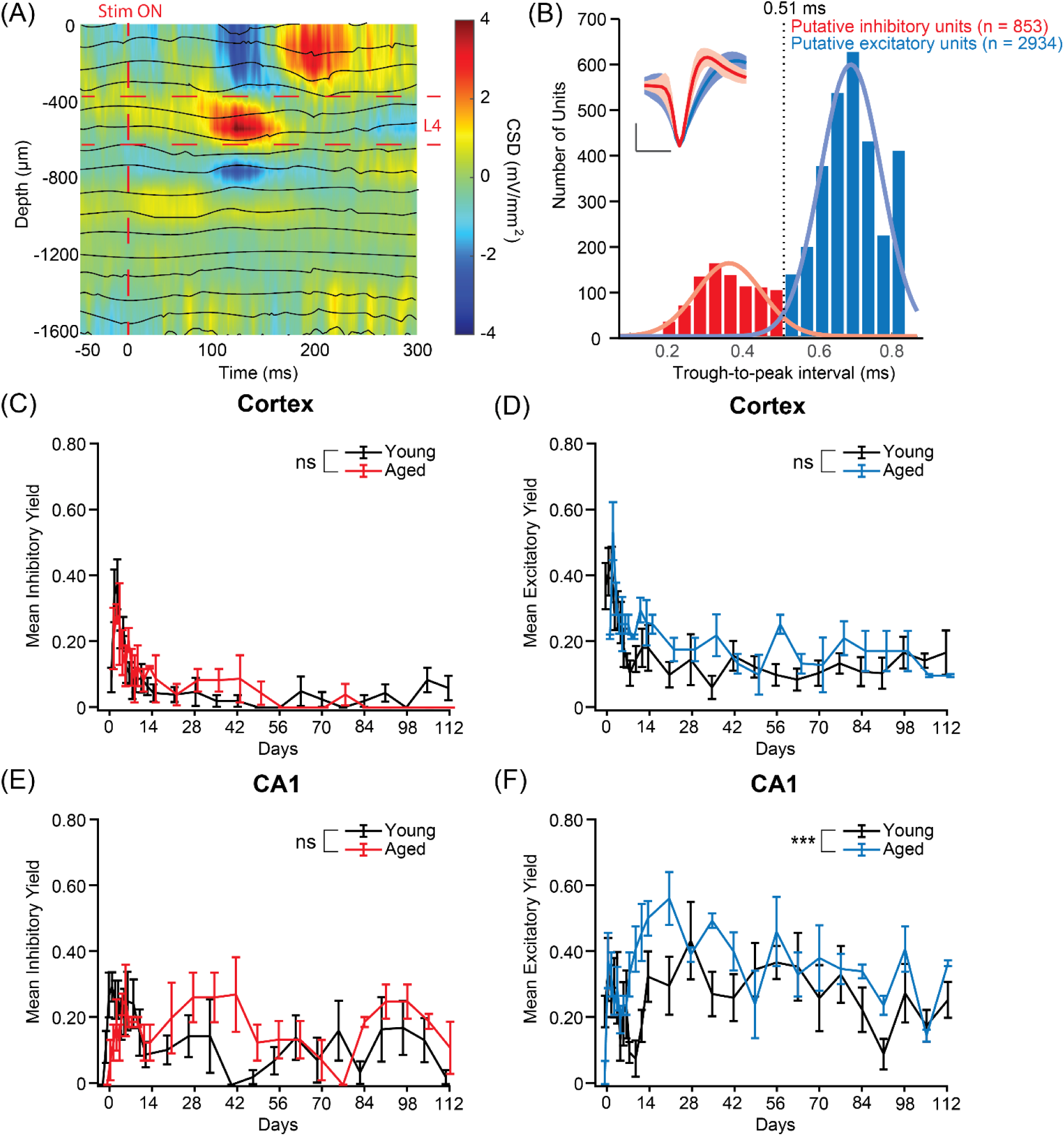
Elevated detection of excitatory activity in CA1 but not in cortical regions. (A) Representative current source density (CSD) heatmap aligned with local field potential (LFP) showing 300 ms of electrical current activity across electrode implant depth after the onset of a visual stimulus drifting gradient. (B) Distribution of spiking activity trough-to-peak intervals (TPI) for classifying putative inhibitory (red, TPI ≤ 0.51 ms and excitatory units (blue, TPI > 0.51). Inset shows representative spiking waveforms (mean ± 1 SD) for each classification. Scale bar represents 0.4ms in time and 0.4uV in amplitude. (C) Inhibitory yield over time in cortex (D) Excitatory yield over time in cortex (E) Inhibitory yield over time in CA1 subregion (F) Excitatory yield over time in CA1 subregion showed significant increases in aged groups. Symbol *** indicates p < 0.001 main group effect determined by linear mixed effects model between young (N = 6 males) and aged (N = 2 males, 4 females) mice. Non-significant differences are indicated by ns.

More interestingly, in the CA1 subregion, the average inhibitory yield detected was slightly elevated but insignificant across time for both groups, but excitatory yield was significantly higher in aged animals (Figure 2E-F, p < 0.05, likelihood ratio test). This is consistent with the postulation that neuron hyperexcitability without a compensatory increase in inhibitory activity occurs throughout aging and age-related disease states (Burke & Barnes, 2006; Menon et al., 2020; Small et al., 2011; Targa Dias Anastacio et al., 2022; Wilson et al., 2005).

### Sensory-evoked MU firing patterns are preserved with age

To evaluate sensory-evoked functional responsiveness at the population level, we presented a visual drifting grating stimulus and recorded multi-unit (MU) activity across V1 and CA1. As SU activity reflects individual neuron firing patterns, MU activity represents population-level responses. As previously published (Chen, Cambi, et al., 2023; Moore et al., 2020; Steven M Wellman et al., 2020), we calculated MU yield as the percentage of electrode site with a significant change in firing rate from the stimulation period compared to the pre-stimulation period (p < 0.05, paired t-test). SNFRR was defined as the magnitude of this change per channel. We assessed MU yield and SNFRR with varying post-stimulus bin sizes and latencies at 1ms resolutions. The temporal responses of MU yield spanning both cortex and CA1 areas were visualized as heatmaps in both young and aged animals (Figure 3A). To compare differences in response latency and SNFRR between groups, we optimized MU yield in young WT mice using bin sizes of 46 ms in cortex and 97 ms in CA1 as previously determined (Chen, Cambi, et al., 2023). Interestingly, we saw no differences in latency and |SNFRR| between young and aged mice (Figure 3B-C), possibly indicating that ensemble evoked firing activity is not impacted by age.

**Figure 3.**
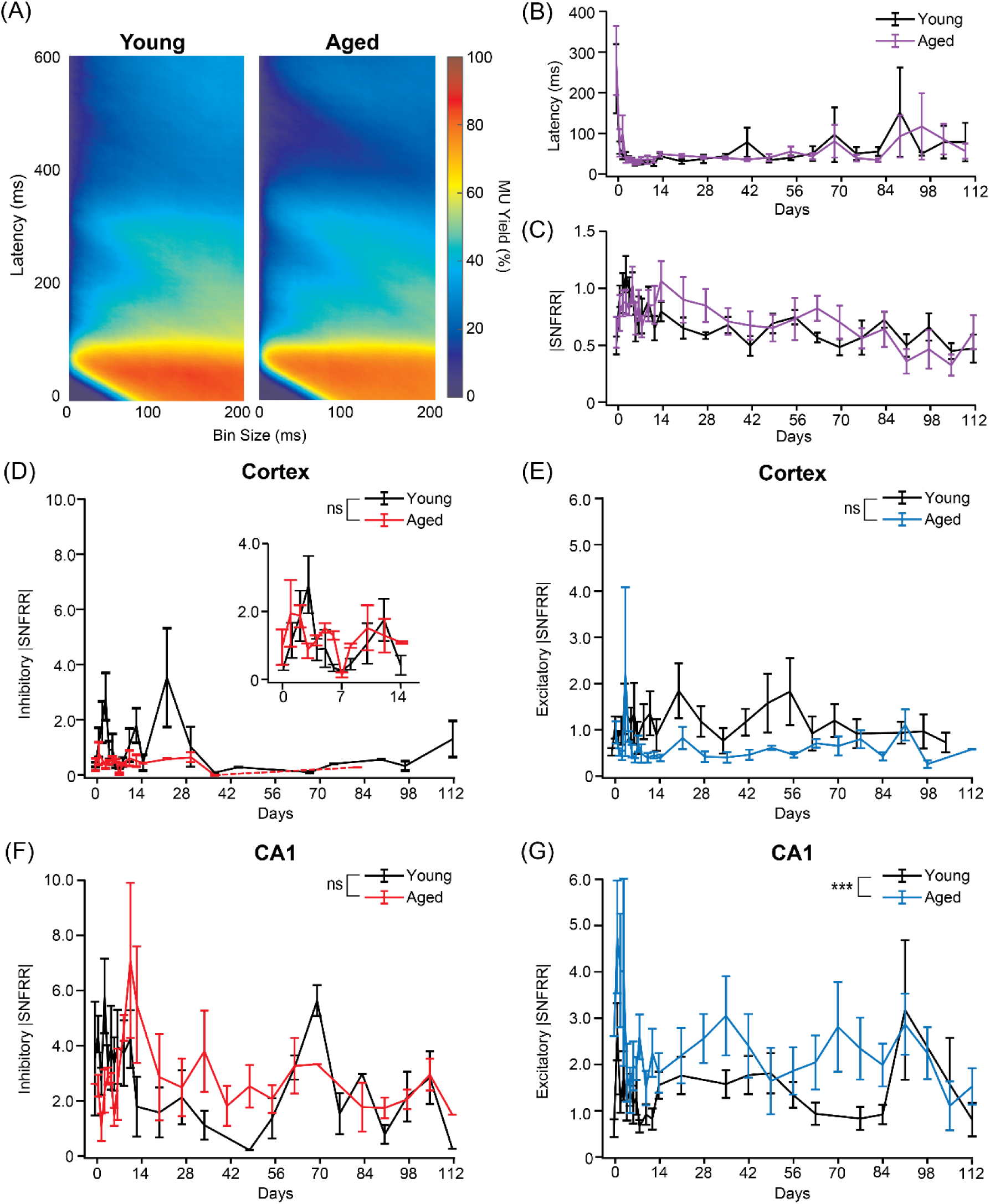
Aging elevates excitatory population-level evoked firing activity in CA1. (A) Mean multi-unit (MU) yield along across all electrode sites plotted as a function of latency and bin sizes at 1 ms resolution following 1 s of visual stimulation for young and aged mice. (B) Mean latency and (C) mean |SNFRR| calculated at the optimal MU yield in young mice and plotted as a function of time for young (black solid) versus aged mice (purple solid). (D) Average |SNFRR| of inhibitory units in the cortex over time for young (black solid) and aged mice (red solid). Inset highlights data from day 0 to day 14. (E) Average |SNFRR| of excitatory units in the cortex over time young (black solid) and aged mice (blue solid). (F) Average |SNFRR| of inhibitory units in CA1 over time for young (black solid) aged mice (red solid). (G) Average |SNFRR| of excitatory units in CA1 was significantly elevated (p = 3.94e-05) in aged mice (blue solid) compared to young mice (black solid). Symbol *** indicates p < 0.001 main group effect determined by linear mixed effects model between young (N = 6 males) and aged (N = 2 males, 4 females) mice. Non-significant differences are indicated by ns.

We then asked whether excitatory-inhibitory imbalances during aging contribute to sensory-evoked responsiveness across regions. As we saw slight increases in inhibitory yield and significant differences in excitatory yield within CA1 over time (Figure 2E-F), we hypothesized that functional activity in CA1 neuronal ensembles would be elevated in the aged group. We found that the average inhibitory and excitatory |SNFRR| in the cortex was trending slightly lower in aged versus young mice (Figure 3D-E), possibly indicating that aging does not largely impact population responsiveness in V1. Again, we found elevated inhibitory |SNFRR| but insignificant between young and aged animals (Figure 3F). However, excitatory |SNFRR| was significantly higher in aged animals compared to young animals (p = 3.94e-05, likelihood ratio test, Figure 3G). Together, these results suggest that there is negligible delay in neural transmission and perserved stimulus responsiveness during normal, healthy aging. Moreover, age-related deficits in behavioral tasks and neural plasticity may be linked to other downstream visuomotor processing areas like the frontal eye field or superior colliculus (Li et al., 2021; Munoz et al., 1998; Young & Hollands, 2012)

### Global oscillatory power declines with age but is not frequency-specific

To evaluate the impact of age on sensory-evoked neuronal oscillatory activity, we analyzed the power spectral density of local field potential (LFP) recorded in young and aged WT mice. Power was computed by applying a base-10 logarithmic transformation to the LFP. The spontaneous power was subtracted from the evoked power to obtain the evoked power across 0.5-125Hz frequencies. Visualizing the normalized power (dB) across implantation depth and the 16-week recording period for young and aged animals (Figure 4A), we observed a notable reduction in power in aged animals across depth and time compared to young mice. Averaging across electrode depth, we found significant reduction in mean power for aged animals compared to young animals (Figure 4B, p < 0.05, likelihood ratio test). To further assess differences is oscillatory activity, we separated the LFP stream to relative power frequency bands: delta (2-4Hz), alpha (7-14Hz), beta (15-30Hz), gamma (30-90Hz), and higher frequency oscillations (HFO, >90Hz). Interestingly, we observed no significant differences for any of the relative frequency bands (Figure 4C-G). Although insignificant, the reduction in relative power within the delta band (Figure 4C) may have contributed more to the overall decline in power.Taken together, these results suggest that aging does not result in a frequency-specific decline in visually evoked oscillatory activity but results in a global reduction that varies with depth. The global reduction potentially indicates impaired network synchrony or neuron responsiveness in the aged cortex.

**Figure 4.**
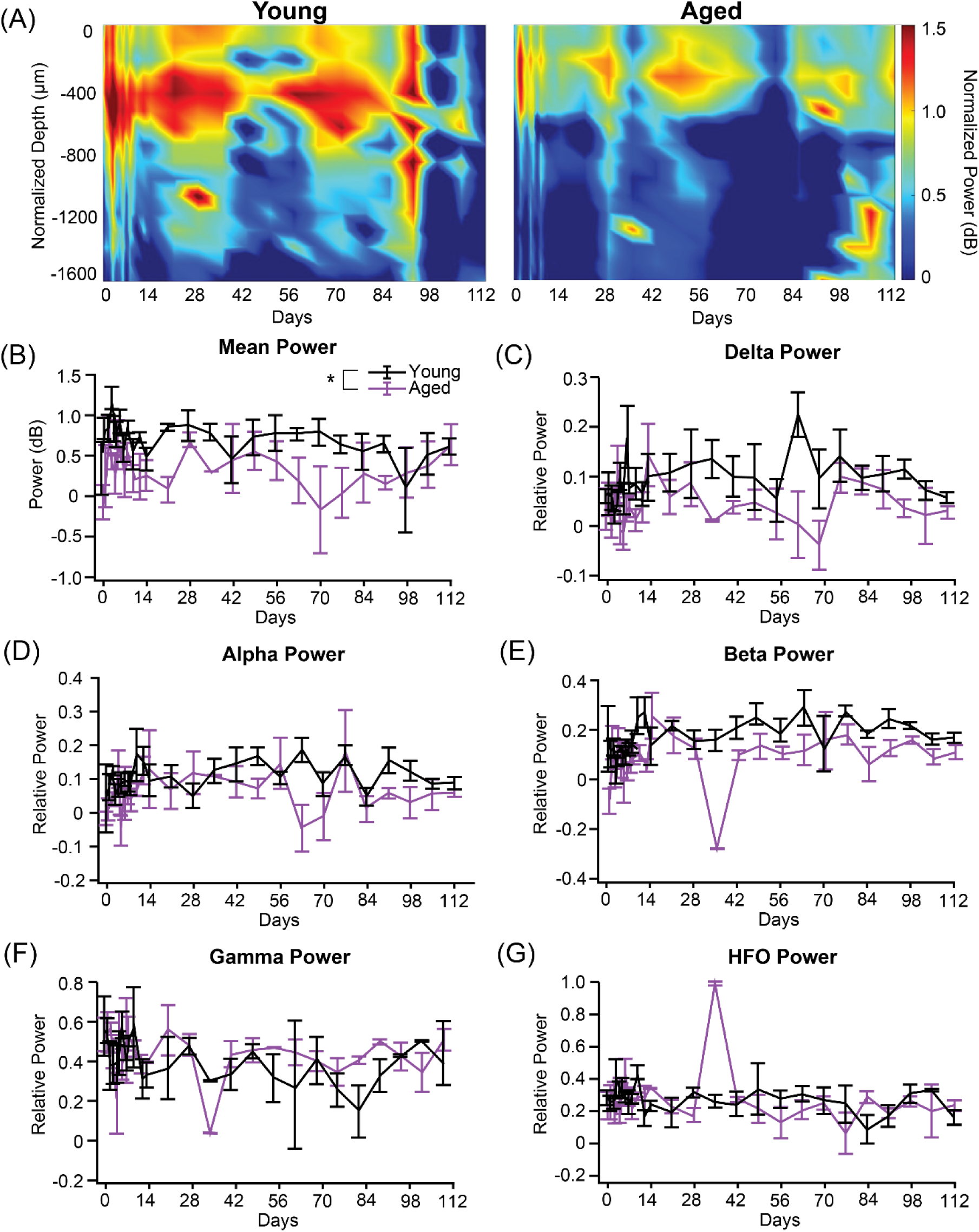
Local field potential power globally diminishes with age in a non-frequency specific manner. (A) Heatmaps of the average normalized visually evoked power plotted as a function of aligned cortical depth and time for young and aged mice. (B) Mean power was significantly reduced in aged mice (purple solid) over time compared to young mice (black solid). Relative power for (C) delta (2-4Hz) (D) alpha (7-14Hz) (E) beta (15-30Hz) (F) gamma (30-90Hz) (G) and high frequency oscillations (HFO, >90Hz) were comparable between groups. This indicates that aging globally reduces that is not frequency specific. Symbol * indicates p < 0.05 main group effect determined by linear mixed effects model between young (N = 6 males) and aged (N = 2 males, 4 females) mice.

### Age Selectively Disrupts Frequency- and Layer-Specific Functional Connectivity

Although diminished neuronal density is a notable feature in the aging brain, we found no significant differences in SU activity (Figure 1) but did observe region-dependent differences in excitatory yield (Figure 2E) and global reduction in LFP mean power (Figure 4), potentially highlighting age-related synaptic discoordination in the neural network. To further investigate network synchrony, we asked whether functional laminar connections were impaired following sensory stimulation in young and aged mice. We used coherence to evaluate the synchronicity in LFP activity throughout the cortex and CA1 regions. Changes in coherence (Δcoherence) were calculated as the difference between the visually evoked and spontaneous cross-spectrum LFP activity across select frequency bands and depths.

We first analyzed Δcoherence from layer IV to layers I and II/III pathways. Encapsulating layer I to layer IV (∼0-500 µm in depth), the frequency-dependent changes in this connection were most prominent on day 5 (Figure 5A), with increases seen in delta and alpha-beta ranges. We saw a main group effect of age for Δcoherence in delta (p = 1.13e-5, likelihood ratio test, Figure 5B) and alpha-beta (p = 1.24e-11, likelihood ratio test, Figure 5C) over time, possibly indicating that aging selectively impairs low- and mid-frequency network synchronization while sparing higher-frequency oscillatory coupling. Significant effects of time were observed in both the alpha-beta (p = 0.0024, likelihood ratio test) and gamma frequencies (p = 0.027, likelihood ratio test). This pattern remains consistent with previous studies showing age-related reductions in delta and alpha coherence across cortical and hippocampal circuits, impairing large-scale network integration and cognitive processing (Chong et al., 2019; Geerligs et al., 2014). Although we found a main effect of time within the gamma frequency (p = 0.027, likelihood ratio test), there was not a main group effect of age (p = 0.195, likelihood ratio test) within gamma frequency coherence (Figure 5D). Previous reports show that gamma frequencies, often linked to sensory encoding, are less vulnerable to age-related decline (Ranasinghe et al., 2025; Voytek et al., 2015).

**Figure 5.**
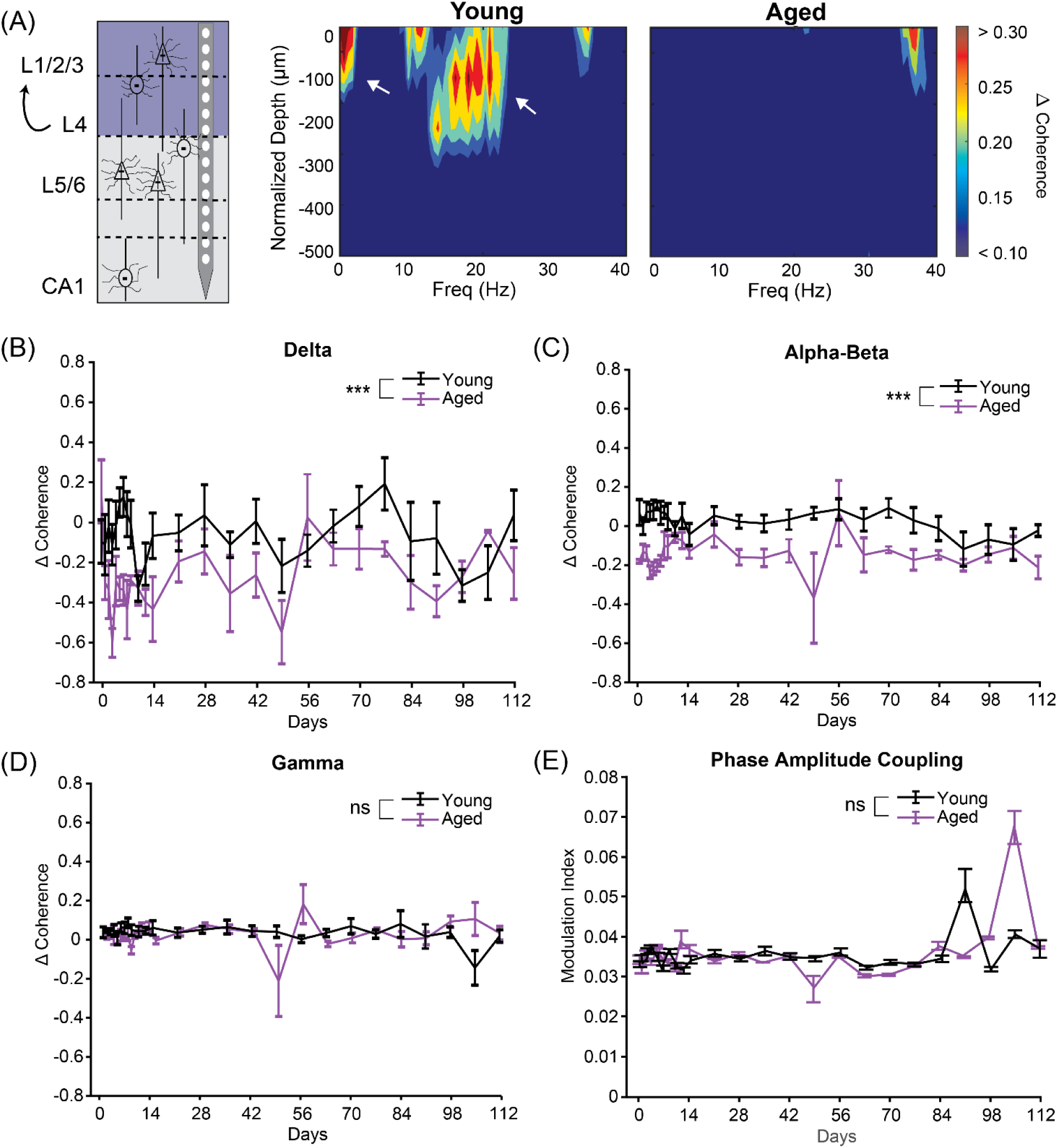
Aging reduces low frequency synchrony of visually evoked integration. (A) Representative graphic highlighting connections from layer IV to superficial layers I and II/III in V1 (left). Heatmaps of visually evoked change in coherence (Δcoherence) as a function of depth (0-500 µm) and frequency for young and aged mice (right). ΔCoherence across (B) delta frequencies showed significantly lower synchrony (p = 1.13e-5) in aged (purple solid) compared to young controls (black solid). (C) Alpha-beta frequencies showed similar significant reductions in aged mice compared to young mice (p = 1.24e-11). (D) Gamma frequency Δcoherence showed no differences between groups (p = 0.027). (E) Quantification of modulation index (MI) between the phase of low-frequency oscillations and amplitude of high-frequency activity indicated no impact of age on functional coordination in layer IV-I/II/III feedforward projection (p = 0.195). The symbol *** indicates p < 0.001 main group effect determined by linear mixed effects model between young (N = 6 males) and aged (N = 2 males, 4 females) mice. Non-significant differences are indicated by ns.

Together, these results suggest that the vulnerability of certain frequency bands to aging may reflect differential reliance of long-range communication compared to local neural communication. To assess functional connectivity between layer IV and layers I and II/III, we calculated the phase amplitude coupling modulation index (MI), where a higher MI indicates stronger coupling. We hypothesized that differences in low frequency bands delta and alpha-beta would result in reduced coupling. However, we saw similar MI values between young and aged groups (Figure 5E), with group and time interaction effect (p < 0.05, likelihood ratio test). These results show that the feedforward interlaminar connections from layer IV to layers I and II/III diminish in temporal coordination without reduced coupling strength, possibly indicating network-level hyperexcitability that is frequency-dependent and region-dependent dysfunction.

Next, we evaluated Δcoherence between superficial (layers I and II/III) and deep (layers V and VI) cortical layers from approximately 200 to 1000 µm. The Δcoherence was most notable on day 77 (Figure 6A). Interestingly, we saw significantly lower Δcoherence between aged and young animals in delta (p = 4.98e-11, likelihood ratio test), alpha-beta (p = 4.00e-9), and gamma (p = 0.033, likelihood ratio test) frequencies (Figure 6B-D). Effects of time were observed in both the alpha-beta (p = 0.026, likelihood ratio test) and gamma frequencies (p = 0.010, likelihood ratio test). These findings align with previous evidence showing that aging disrupts oscillatory synchrony during large-scale cortical processing. Age-related reduction in lower frequency oscillations (delta and alpha bands) has been associated with lower synaptic density and cholinergic input from basal forebrain neurons, which play a major role in network coordination and excitatory-inhibitory balance (Chaves-Coira et al., 2023). Further, impaired oscillatory activity in both low and high frequencies in interlaminar communication is often associated with deficits in sensory integration and memory (Hernandez et al., 2020; Radulescu et al., 2023). The observed decline in coherence across all frequency bands is consistent with findings at both local circuitry and large-scale disruption in network connectivity (Bajaj et al., 2017; Talelli et al., 2008). Again, we found comparable values for MI comparing young and aged groups, with a group and time interaction effect (p < 0.05, likelihood ratio test, Figure 6E). Superficial projections from layer I and II/III towards deeper layers V and VI are critical for information processing. Thus, these results suggest that the integrity of ascending and descending interlaminar communication is particularly vulnerable to age-related disruption of processing visual input towards downstream regions.

**Figure 6.**
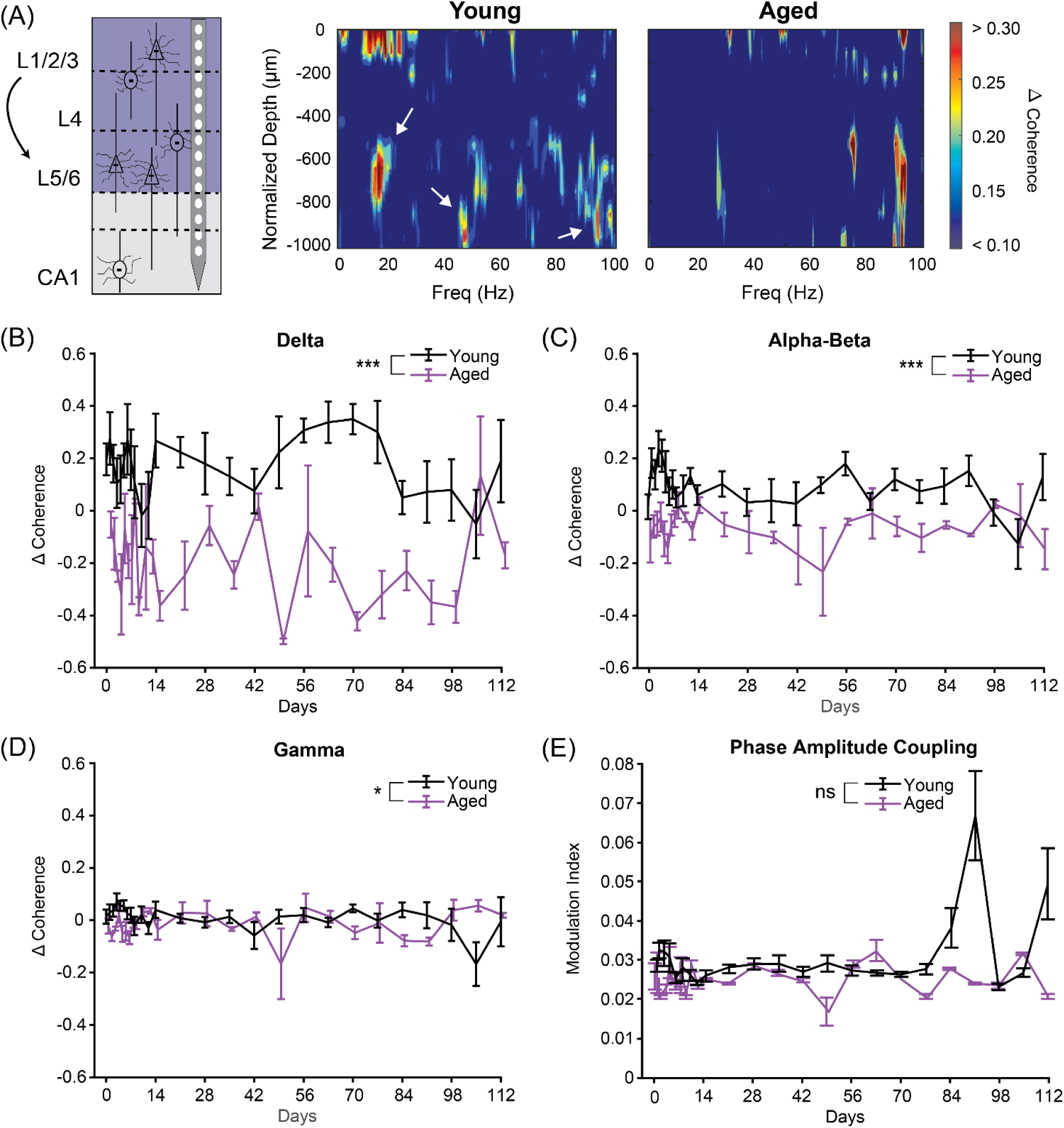
Aging reduces ensemble synchrony for visual input processing across frequency bands. (A) Representative graphic highlighting connections from superficial layers I and II/III to deeper layers V and VI (left). Heatmaps depict Δcoherence as a function of depth and frequency for young and aged mice (right). Significant reductions in Δcoherence across (B) delta (p = 4.98e-11), (C) alpha-beta (p = 4.00e-9), and (D) gamma frequencies (p = 0.033) were observed in aged (purple solid) compared to young mice (black solid). (E) Quantification of MI showed comparable trends between young and aged mice. Respectively, * and *** indicate p < 0.05 and p < 0.001 main group effect determined by linear mixed effects model between young (N = 6 males) and aged (N = 2 males, 4 females) mice. Non-significant differences are indicated by ns.

Lastly, we analyzed the corticohippocampal projections spanning across layers V and VI to CA1 subregion of the hippocampus (∼400-1600 µm, bottom electrode site). The long-range projections from deep-layer pyramidal neurons are essential for relaying processed sensory information to the hippocampus, facilitating both memory encoding and retrieval (Eichenbaum, 2017; Swanson & Cowan, 1977). Similarly to previous laminar connections, we observed prominent changes in coherence on day 5 (Figure 7A). Once again, we found a main effect of age on Δcoherence within delta frequencies (p = 1.60e-3, Figure 7B). Interestingly, Δcoherence were comparable between young and aged animals within alpha-beta (p = 0.276, Figure 7C) and gamma bands (p = 0.816, Figure 7D). We found no effect of time were observed across alpha-beta (p = 0.885), delta (p = 0.713), or gamma frequencies (p = 0.867). This selective vulnerability of delta coherence is consistent with previous findings that low-frequency oscillations are critical for coordinating large-scale communication between cortical and hippocampal regions, particularly during memory consolidation and retrieval (Benchenane et al., 2010; Sirota et al., 2008). Age-related reductions in delta synchrony have been linked to decreased synaptic integrity and impaired neuromodulatory input, which disproportionately affect long-range projection systems (Beason-Held et al., 2017; Chaves-Coira et al., 2023). The absence of significant age effects in the gamma bands suggests that higher-frequency oscillatory communication, often associated with local processing and fast information transfer, may be preserved in corticohippocampal circuits during normal aging (Başar, 2013; Colgin, 2016). Consistent with other interlaminar connections, we found comparable trends for MI for young versus aged groups, with a group and time interaction effect (p < 0.05, likelihood ratio test, Figure 7E). Taken together, these results indicate that aging specifically disrupts the slow oscillatory synchrony necessary for effective communication between deep cortical layers and the hippocampus, which may underscore age-related deficits in memory and sensory integration.

**Figure 7.**
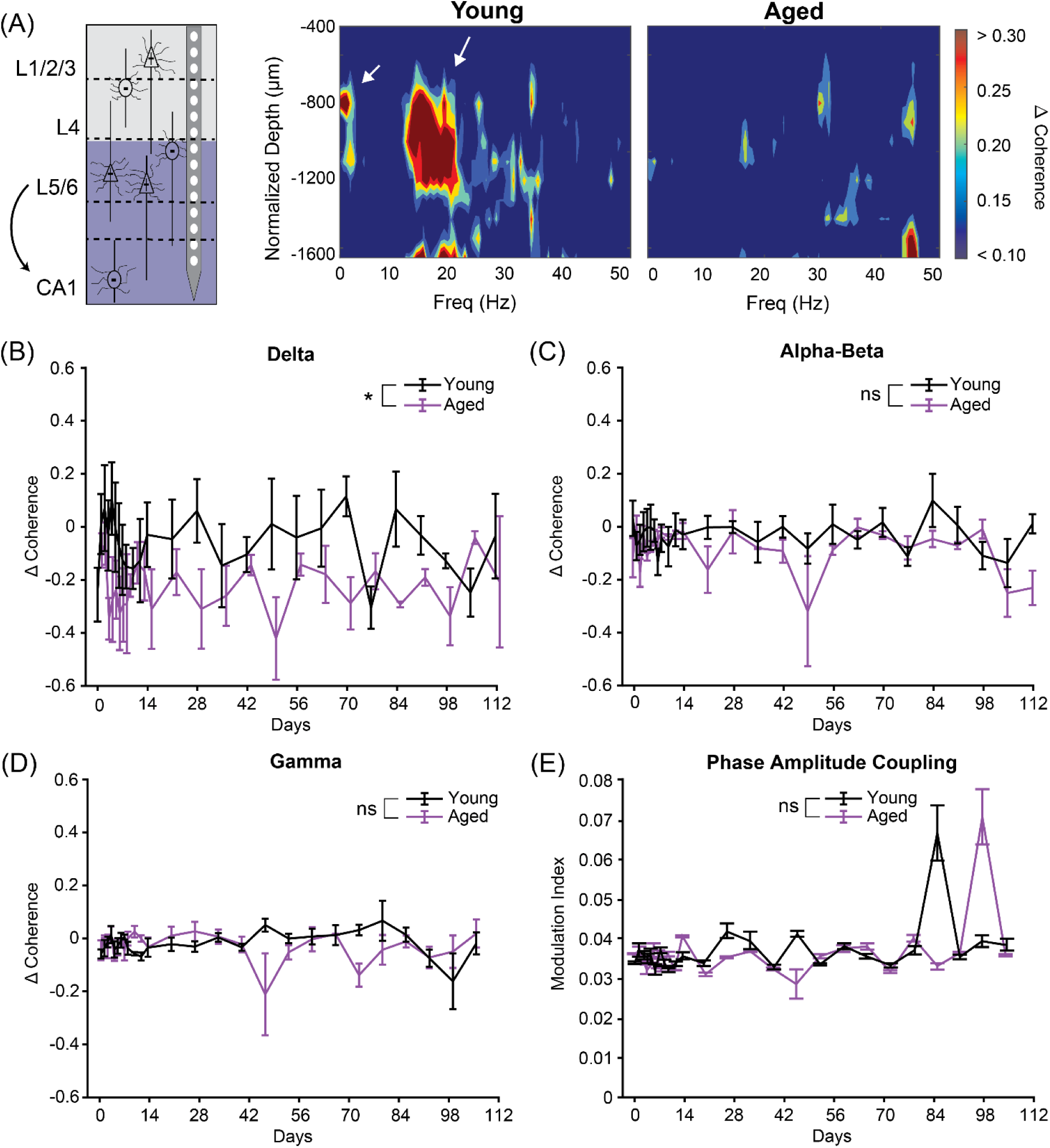
Aging reduces low frequency synchrony projections from V1 to CA1. (A) Representative graphic highlighting projections from layers V/VI from V1 towards CA1 subregion of the hippocampus (left). Heatmaps depict Δcoherence as a function of depth and frequency for young and aged mice (right). (B) Significant reductions in Δcoherence (p = 1.60e-3) across delta frequencies was observed in aged mice (purple solid) compared to young controls (black solid). ΔCoherence values were insignificant across (C) alpha-beta (p = 0.276) and (D) gamma frequencies (p = 0.816) between groups. (E) Quantification of MI showed comparable trends between young and aged mice. Symbol * indicates p < 0.05 main group effect determined by linear mixed effects model between young (N = 6 males) and aged (N = 2 males, 4 females) mice. Non-significant differences are indicated by ns.

## DISCUSSION

The central theme of this study distinguishes single-neuron firing patterns from population-level changes in functional connectivity. We found preservation of sensory-evoked single-unit (SU) activity in chronically implanted mice across the rodent lifespan, from young adult (2 months) to middle-to-late adulthood (13 months). Employing a drifting grating visual stimulus, we saw frequency-selective desynchronization in neuronal populations without significant alterations to cross-frequency interactions across laminae and between brain regions.

### Aging Does Not Impair the Reliability of Chronic Neural Recordings

Age-related physiological changes, such as dendritic spine loss and glial morphology, can degrade the robustness of chronic electrophysiology. Dendritic spines, the primary postsynaptic sites for excitatory synapses, decline 20-40% in cortical and hippocampal regions over a lifetime, weakening synaptic connectivity and plasticity (Dumitriu et al., 2010; Feldman & Dowd, 1975; Mrak et al., 1997). In parallel, glial cells can adopt a senescent phenotype and promote pro-inflammatory pathways and destabilize the synaptic microenvironment (Hawkley & Cacioppo, 2004; Shafqat et al., 2023). More crucially, these age-related structural and glial cell changes correlate with cognitive deficits in aged preclinical models and humans, impairing both spatial and working memory (Burke & Barnes, 2006; Morrison & Baxter, 2012). However, most preclinical studies focus on young animals, whereas clinical devices target older adults for lifelong implantation (Chestek et al., 2011; Flesher et al., 2021; Losanno et al., 2023). Thus, investigating longitudinal electrophysiological impairments is critical for the translatability and understanding of the aging brain.

In addition to aging itself, concerns over implant-induced immune responses have further limited longitudinal in vivo studies. Foreign body responses, including local increases in reactive oxygen species (ROS) and glial reactivity, complicate the ability to separate device-related effects from aging effects (Bardaweel et al., 2018; Li et al., 2023; Liguori et al., 2018). The accumulation of ROS leads to heightened oxidative stress that can impair normal supportive functions in the tripartite synapse (Rabah et al., 2025). Aged astrocytes exhibit elevated intracellular ROS and calcium overload, which activate stress- and cell-death related signaling paths such as JNK/SAPK (Ishii et al., 2017). Accordingly, these deficits can cause these glial cells to shift toward a reactive, pro-inflammatory phenotype and secrete cytokines and chemokines that destabilize the synaptic environment (Boisvert et al., 2018). Neurons are particularly vulnerable to ROS-induced damage, leading to metabolic dysfunction and weakened synaptic connectivity (Bellaver et al., 2017). Elevated oxidative stress can lead to DNA damage, protein oxidation, and energetic deficits that accelerate cell senescence and susceptibility to apoptosis (Bezprozvanny & Mattson, 2008). Across a lifespan, dysfunctional astrocytes and damaged neurons exacerbate ROS production and continuously drive progressive cognitive decline and neurodegeneration in aging (van Deijk et al., 2017). Thus, few studies have elected to utilize chronic in vivo recordings due to concerns over implant-induced immune responses.

Despite aging and implant-induced changes, chronic extracellular recordings remain robust in aged mice. Although some studies have shown that aging does not alter SU activity in the motor cortex (Nolta et al., 2024) as well as in the entorhinal cortex (Merrill et al., 2000), it remains unclear whether this observation is consistent across regions and occurs simultaneously. We found that SU yield (Figure 1B-C), SNR (Figure 1D), and SNR per active site (Figure 1E) were unaffected by age, confirming that longitudinal electrophysiological recordings in aged animals are reliable for assessing network-level changes. Thus, age-related declines in oscillatory power (Figure 4), cross-frequency coupling, and functional connectivity (Figures 5-7) may result from physiological mechanisms rather than recording artifacts or progressive glial scarring.

### Age-Related Hyperexcitability in CA1 Circuits Were Not Observed in V1

We found hyperexcitability in aged CA1 neurons, while cortical neurons in V1 did not show this, consistent with prior studies indicating that memory-impaired aged rodents exhibit heightened neural excitability in hippocampal regions, such as elevated cFos expression and hyperactive CA3 neurons (Haberman et al., 2017). Importantly, the cortex maintained a stable balance between excitation and inhibition (Figure 2C-D). In contrast, hyperexcitability in CA1 (Figure 2E-F) appeared to arise from a disrupted excitatory-inhibitory balance rather than widespread neuronal loss, highlighting the hippocampus as a selective target of age-related circuit dysfunction. To capture how these cellular changes scale to the network level, we assessed ensemble spiking activity, revealing stronger excitatory |SNFRR| responses in CA1 (Figure 3G), indicating that aging alters intrinsic cellular activity and amplifies sensory-evoked responses across hippocampal networks. A likely explanation for this difference lies in the intrinsic properties of CA1 pyramidal neurons. Compared to cortical neurons, aged CA1 cells fire more frequently, a change that can be traced to higher input resistance and altered ion channel conductance (Brahimi et al., 2023; Campbell et al., 1996). These biological shifts influence both action potential firing and hyperpolarization periods (Brahimi et al., 2023), mechanistically linking age-dependent ion channel remodeling and network-level hyperexcitability. At the population level, this hyperexcitable state of CA1 neurons is also reflected in the reduced LFP power (Figure 4B), likely due to impaired spike timing and weaker synaptic integration (Peace et al., 2024). This paradoxical combination of heightened single-cell excitability coupled with weakened population synchrony reveals a foundational breakdown in hippocampal signal integration during aging, a process that underlies memory formation and retrieval.

Age-related changes in myelination likely contribute to the selective hyperexcitability observed in CA1 but not in V1. Myelin integrity is essential for maintaining rapid and precise neural signal conduction, critical for synchronizing hippocampal network activity (Ahn et al., 2017). Studies in aging rodents and primates have shown that hippocampal subregions experience significant reductions in myelin basic protein (MBP) expression, myelin integrity, fragmentation and thinning compared to the cortex (Peters, 2009). These myelin deficits slow conduction velocity and degrade temporal coordination, promoting network hyperexcitability (Murray et al., 2023). Conversely, the cortex may maintain or remodel myelin more effectively during aging, sustaining stable excitatory/inhibitory balance and preventing aberrant firing patterns (Shepherd et al., 2012). Additionally, myelin loss in the hippocampus is linked with reactive glial activation and metabolic stress, factors which further exacerbate circuit instability (Ahn et al., 2017). In some cortical areas, myelination continues or is enhanced with age through the addition of lamellae and remyelination by oligodendrocytes (Hill et al., 2018; Liu et al., 2025), potentially preserving the fidelity of excitatory-inhibitory balance and preventing aberrant neuronal hyperactivity. This preservation of cortical myelin integrity may underline the absence of hyperexcitability in aged V1 despite widespread age-related neural changes elsewhere. Together, these findings suggest that hippocampal hyperexcitability emerges from the convergence of intrinsic membrane remodeling and region-specific myelin vulnerability, while preserved cortical myelination and excitatory-inhibitory balance protect V1 from similar dysfunction.

### Aging is Associated with Frequency-Specific Laminar Desynchronization

Regional and frequency-dependent laminar desynchronization was evident in the aged mice compared to young controls. Although global LFP power declined (Figure 4B) without selective loss in any frequency band (Figure 4C-G), population-level analysis revealed pronounced desynchronization of low-frequency delta and alpha-beta bands in V1-CA1 pathways (Figures 5-7). Importantly, connectivity changes were not uniform, with all frequency bands disrupted between superficial cortical layers I and II/III and deeper layers V/VI (Figure 6B-D), and selective low-frequency fluctuations in other interregional connections (Figure 5B-D and 7B-D). These results suggest that temporal coordination is a critical network feature disrupted by aging, rather than simple power loss and despite preserved individual neuronal firing. Laminar cross-frequency coupling, however, remained largely intact (Figures 5E, 6E, and 7E), suggesting that local phase-amplitude interactions persist even when long-range timing is impaired.

In chronically implanted neural interfaces, these age-related vulnerabilities can be compounded by device-tissue interactions (Sridharan et al., 2013). Chronic implantation of these devices compromises myelin integrity, leading to acute demyelination, myelinosome formation, and abnormal tau phosphorylation that impairs oligodendrocyte-astrocyte coordination and metabolic support (Chen et al., 2021; Gupta et al., 2024; Stice et al., 2007). Because myelin integrity is essential for rapid conduction velocity and temporal precision (Suminaite et al., 2019), such impairments exacerbate age-related myelin loss and are likely contributing to interlaminar desynchronization along V1-CA1 pathways (Figures 5-7). Pro-myelinating Clemastine has been previously shown to improve SU recording quality, while a demyelinating cuprizone model reduced (Chen, Cambi, et al., 2023; Steven M Wellman et al., 2020). Thus, myelin maintenance emerges as a critical determinant of both functional connectivity and signal propagation. These neuroinflammatory processes may differentially impact layers with distinct glial densities and contacts, thereby contributing to layer-specific desynchronization seen with aging.

Notably, recent computational and experimental work indicates that partial remyelination can restore conduction velocity and improve circuit timing, while modulating astrocyte reactivity may rescue synaptic balance and oscillatory coherence (Dimovasili et al., 2023).

Metabolic dysregulation and impaired neurovascular coupling (NVC), linking neurons, astrocytes, and cerebral blood flow, disrupts oscillatory coordination. Aging impairs NVC, reducing delivery of oxygen and nutrients, increasing oxidative stress, limiting ATP production, and heightening the risk of cognitive decline (Shih et al., 2013; Snyder et al., 2015; Yabluchanskiy et al., 2020). As a result, cellular energetics and sustained oscillatory activity worsen over time. With chronic implantation of microelectrodes, these effects become exacerbated by creating focal ischemic environment around the injury site (Wellman et al., 2023). Astrocytic dysfunction further compounds these effects by altering the extracellular milieu and synaptic homeostasis, weakening NVC, and disrupting ion balance essential for generating and maintaining LFPs across cortical layers (Stogsdill et al., 2023). Because astrocyte density and vascular contacts vary by layer (Endo et al., 2022), layer-specific astrocytic and vascular vulnerabilities likely contribute to the differential desynchronization observed in aging. Notably, computational and experimental studies suggest that partial remyelination can restore conduction velocity and improve circuit timing, while modulating astrocyte reactivity may help rescue synaptic balance and oscillatory coherence (Dimovasili et al., 2023). Together, these deficits suggest that aging yields two parallel distinctions: preserved local cross-frequency interactions but impaired long-range laminar and interregional coordination. The degradation of LFP temporal alignment (Figures 4B, 5B-D, 6B-D, 7B-D), despite stable SU activity (Figure 1C-E), illustrates that neuronal firing reliability is not sufficient to maintain population-level synchrony. This dissociation between intact single-neuron activity and disrupted network coherence underscores the importance of examining both cellular and circuit scales when assessing brain aging.

Finally, our findings demonstrate that aging does not impair the reliability of chronic extracellular recordings, as SU yield and SNR remained stable across the rodent lifespan. Instead, aging selectively alters circuit dynamics, with hippocampal CA1 hyperexcitability absent in V1, and frequency-specific laminar desynchronization weakening long-range coordination while sparing local cross-frequency interactions (Figures 5-7). This selective vulnerability highlights chronic recordings as powerful tools to identify region- and frequency-specific network changes that may represent early signs of cognitive decline.

### Limitations, future directions, and conclusion

A limitation of the current study is the number of electrode sites residing in V1 and CA1. Although the single-shank multi-channel microelectrode used in this study allowed for assessment of laminar dysfunction and degradation of the neuronal population, the electrode site density positioned in each layer and between cortex and CA1 areas were sparse. Further, our microelectrodes failed to detect putative inhibitory waveforms in the cortex past day 84 (Figure 2C). This may be a limitation of the electrode site radius. While this may partly reflect geometric limitations of the recording sites, which capture activity within an approximate 50 µm radius depending on implantation location (Buzsáki, 2004), inhibitory activity was readily observed earlier in the implantation period. This pattern suggests that the later absence of inhibitory signals may reflect selective vulnerability of inhibitory neurons or their synaptic integration under chronic implant and aging conditions, rather than being solely attributable to electrode location (Atkinson et al., 2021; Gupta et al., 2024; Salatino et al., 2017). Future studies may address these limitations with electrode designs of higher recording site density, such as Neuropixels or advanced microprobes, capable of spanning larger regions of visual cortex and their projections into the CA1 subregion of the hippocampus (van Beest et al., 2025). Implanting higher resolution and higher density electrodes to probe the neural activity will elucidate region-specific circuitry changes and excitatory-inhibitory imbalances due to aging. Increased recording channel density will allow for a better assessment of population-level neuronal desynchronization and degrading spiking activity in the aged brain. Due to availability and survival of animals during surgical procedures, this study compares electrophysiological differences in 6 young male mice versus 2 aged male and 4 aged female mice. Future studies will address how sex-differences may impact the neural network and its functional laminar connectivity.

In summary, chronic electrophysiological recordings reveal that sensory-evoked single-unit activity remains largely preserved across the rodent lifespan, while functional laminar and cortico-hippocampal connectivity shows depth-dependent degradation. Disruptions in layer II/III interlaminar connectivity and modulated CA1 spiking activity disrupts slow oscillatory synchrony, indicating network-level vulnerabilities that distinguish normal aging from pathological neuronal loss. Local cross-frequency coupling remains intact, indicating that individual neuronal populations retain functional capacity despite impaired population-level temporal coordination. Laminar desynchronization and disrupted long-range timing likely arise from age-related myelin and synaptic degradation, neurovascular uncoupling and metabolic stress, oxidative stress, neuroinflammation, and cellular senescence. These findings are essential for directly addressing the shortcomings of current chronic neural interfaces for older individuals, informing future strategies aimed at preserving excitatory-inhibitory balance, maintaining myelin integrity, enhancing neurovascular coupling, and minimizing neuroinflammation for device longevity.

## DATA AVAILABILITY

The source data are available to verified researchers upon request by contacting the corresponding author.

## ACKNOWLEDGMENTS

This work was supported by: NIH R21NS108098, NIH R01NS105691, NIH R01NS115707, NIH R03AG072218, NIH R01NS129632, F99NS124186, and NSF CBET CAREER 1943906. The authors would like to thank Kevin Stieger and Keying Chen for helpful discussions and guidance with multi-unit activity and phase-amplitude coupling analyses as well as Adam M. Forrest for valuable feedback throughout the writing process.

Present address of S.M.W.: New York City, New York, United States. Present address of N.S.: Kyoto, Japan. Present address of S.S.: San Francisco, California, United States. Preprint is available at https://doi.org/.

## DISCLAIMERS

The authors have no disclaimers.

## AUTHOR CONTRIBUTIONS

S.M.W., F.C., T.D.Y.K., conceived and designed research; S.M.W., S.S., performed experiments; T.T.D.T, S.M.W., N.S., C.G.P., analyzed data; T.T.D.T, S.M.W., interpreted results of experiments; T.T.D.T, S.M.W., N.S., C.G.P., prepared figures; T.T.D.T drafted manuscript; T.T.D.T, S.M.W., N.S., C.G.P., T.T., F.C., T.D.Y.K., edited and revised manuscript; T.T.D.T, S.M.W., N.S., C.G.P, T.T., F.C., T.D.Y.K., approved final version of manuscript

